# OxLDL/LOX-1 Mediated Sex, Age, and Cell Dependent Alterations in Mouse Thoracic Aortic Vascular Reactivity

**DOI:** 10.1101/2023.09.07.556764

**Authors:** Trevor S. Wendt, Rayna J. Gonzales

**Affiliations:** Department of Basic Medical Sciences, University of Arizona, Phoenix, Arizona, USA

**Keywords:** Endothelium, Oxidized Low-Density Lipoprotein, Sex Differences, Thoracic Aortic Stiffness, Vasoreactivity

## Abstract

Elevated oxidized low-density lipoprotein (oxLDL) is a risk factor and component that worsens cardiovascular disease states. OxLDL can elicit its detrimental action, via lectin-like oxLDL receptor 1 (LOX-1) and has been shown to disrupt vascular function. In this study, we determined whether oxLDL, via LOX-1, alters aortic vascular reactivity and determined if sex differences exist. Thoracic aortic endothelium-intact or -denuded ring segments were isolated from intact C57BL/6J female and male mice and incubated with oxLDL ex vivo (50ug/dL; 2h). Using wire myography, cumulative concentration-response curves to phenylephrine (PE) were generated to determine contractile responses. From these curves, the EC50 was determined and used to contract rings to assess acetylcholine (ACh) dependent relaxation. Calculated aortic stiffness and remodeling, as well as mRNA expression of vasoactive and pro-inflammatory mediators were assessed. BI-0115 (10*μ*M; selective LOX-1 inhibitor) was used to determine LOX-1 dependence. We observed differential sex, age, endothelial cell, and LOX-1 dependent alterations to the efficacy of PE-induced contractile responses and ACh-mediated vasorelaxation in the thoracic aortic rings following oxLDL exposure. Additionally, we observed a distinct sex and age effect on thoracic aortic stiffness following exposure to oxLDL. There was also a sex effect on calculated vessel diameter, as well as an age effect on oxLDL-mediated inward remodeling that was LOX-1 dependent. Thus, LOX-1 inhibition and the resulting attenuation of oxLDL/endothelial-mediated alterations in aortic function suggests that there are differential sex differences in the role of oxLDL/LOX-1 in the thoracic aorta of male and female mice.

**NEW & NOTEWORTHY:** We investigated the effects of oxidized low-density lipoprotein (oxLDL) via the LOX-1 receptor on murine thoracic aortic vasoreactivity, stiffness, and remodeling across age and sex. Acute exposure to oxLDL led to altered vasoreactivity, endothelial dysfunction, and changes in aortic stiffness and remodeling. These effects were in-part age, sex, endothelial, and LOX-1 dependent. This study reveals potential complex interactions in oxLDL/LOX-1-mediated vascular responses that could serve as potential therapeutic intervention for vascular diseases such as atherosclerosis.

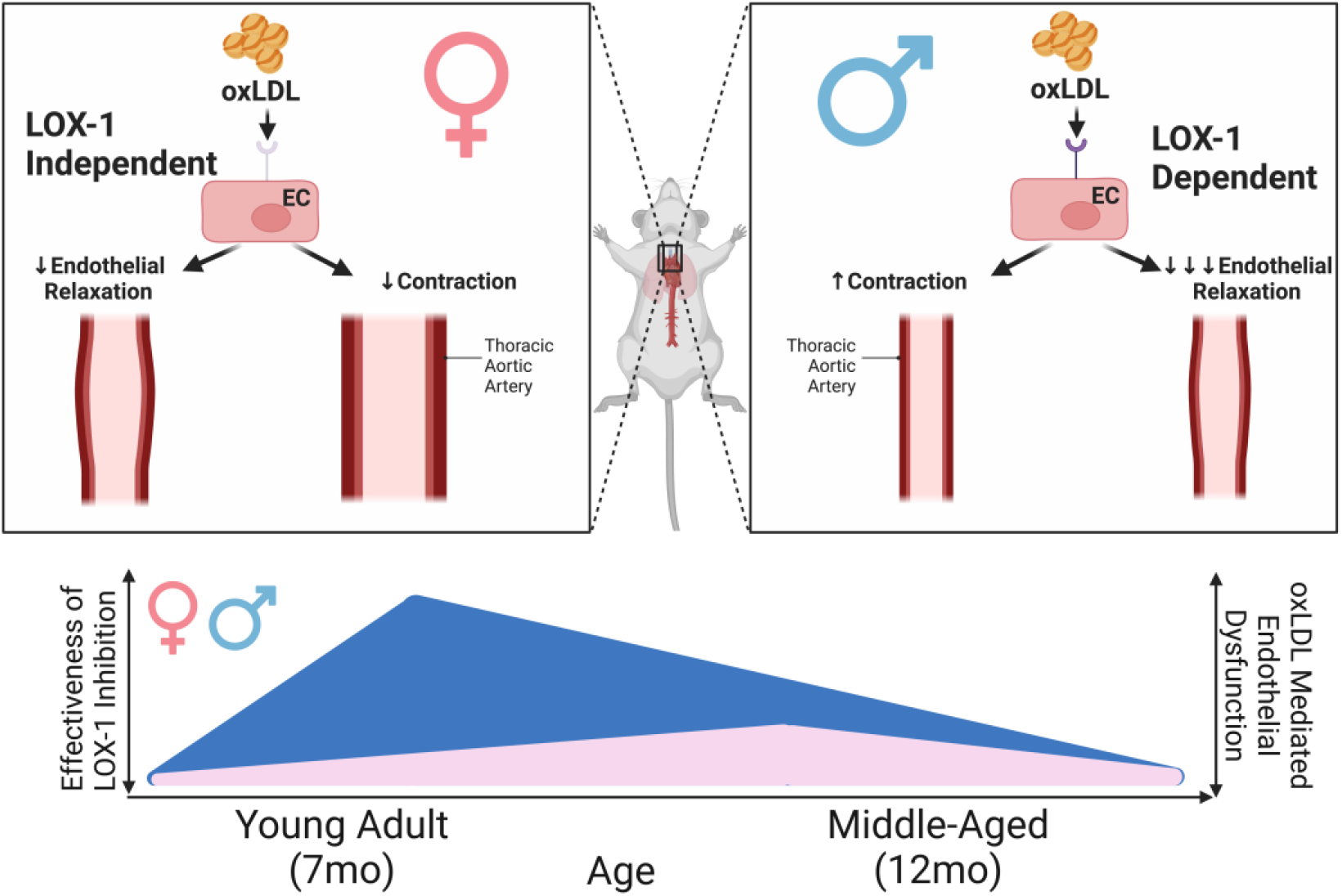

## INTRODUCTION

Elevated oxidized low-density lipoprotein (oxLDL) is a risk factor and key component in cardiovascular disease pathogenesis. The increasing prevalence of dyslipidemia not only confined to the elderly (1) has sparked interest into the molecular and functional consequences of oxLDL acting on the vasculature. Coupled with a rise in metabolic syndromes such as obesity and diabetes mellitus as well as cardiovascular diseases including atherosclerosis (2) which has been linked to increased levels of oxLDL (3), indicates the need for further elucidation of the impact of oxLDL on vascular health and function. Previous investigations have provided valuable insights into the impact of oxLDL on the regulation of aortic vasoreactivity, encompassing various aspects such as endothelial dysfunction, nitric oxide-mediated dilation, and redox control; however, to the best of our knowledge there has not been a direct sex comparison in older middle aged (7 mo. and 12 mo.) adult mice.

Lectin-like oxLDL receptor 1 (LOX-1), a type II integral membrane glycoprotein receptor (4), when activated has been shown to disrupt vascular function (5-12) leading to altered vasoreactivity. Sex differences in vasoreactivity have been previously noted (13, 14); however, the effect of sex on oxLDL aortic vascular reactivity via LOX-1 remains an area of investigation. Research into the effects of oxLDL acting through the LOX-1 receptor on the mouse thoracic aorta has yielded valuable insights into various aspects of cardiovascular physiology. One critical area of investigation is vasocontractility and -relaxation, where involvement of LOX-1 receptor via the endothelium plays a pivotal role. LOX-1 receptor activation by oxLDL contributes to endothelin-1-mediated vasoconstriction and impairs vasorelaxation through NO depletion (4) signifying that this pathway might play a multimodal role in regulating vascular tone and reactivity. Additionally, LOX-1 deficiency has been associated with the thinning of adventitial collagen, potentially contributing to increased susceptibility to ruptured abdominal aortic aneurysms (15). These studies suggest that LOX-1 also plays a role in altering structural integrity and highlights the need for further exploration into the mechanisms underlying aortic stiffness and remodeling in response to oxLDL-mediated LOX-1 signaling.

One of the preceding indications of development of vascular pathology such as atherosclerosis in larger vessels is decreased endothelial-dependent vasodilation which can predispose vessels to structural changes (16). It has been previously shown that oxLDL can exhibit direct inhibitory effects on nitric oxide (NO)-mediated vessel dilation, indicating a critical link between oxLDL and endothelial dysfunction (8, 17-19). Furthermore, it has been found that factors including diet, sex, and aging play a role on endothelial dysfunction in the apolipoprotein-E deficient mouse model, suggesting a highly variable interplay between oxLDL, endothelial function, and aortic health (20). The role of oxLDL has been further emphasized in the intricate and multimodal cascade of endothelial dysfunction and contributions to diseases such as atherosclerosis (21). One of the hallmarks of oxLDL-mediated processes and related to endothelial dysfunction is, oxidative stress, which has been a central theme in the study of cardiovascular disease. This is further convoluted by the addition of inflammation which has also been investigated in the context of atherosclerosis and demonstrates the intricate mechanisms through which oxLDL may impact aortic vasoreactivity.

Endothelial dysfunction, during pathology such as atherosclerosis, has also been subject of investigation in the context of oxLDL-LOX-1 interactions. Studies have demonstrated that LOX-1-mediated oxidative stress and inflammation can lead to endothelial dysfunction, a critical step in atherogenesis (22). Moreover, the reciprocal relationship between LOX-1 and adiponectin has been implicated in endothelial dysfunction regulation, emphasizing the complexity of molecular interactions that regulate vascular reactivity (23). Notably, sex differences have been an emerging theme in this area of research (24). Endothelial overexpression of LOX-1 has been found to increase plaque formation and atherosclerosis in male mice (25), which coupled with independent findings of increased LOX-1 expression in murine male aortas compared to females suggests that the impact of LOX-1 signaling may be influenced by sex (26). These previous findings together emphasize the need for further elucidation of the detrimental role oxLDL/LOX-1 plays on vascular wall health and function including the consideration of sex-specific responses.

## MATERIALS AND METHODS

### Mice

All mouse experiments were approved by the Institutional Animal Care and Use Committee at the University of Arizona (IACUC 16-079, PI: RJG). Male and female mice (C57BL/6J; Jackson Laboratory Bar Harbor, ME) were non-apolipoprotein-E deficient mice to minimize confounding variable of prior exposure to elevated serum levels of oxLDL (4.5ng/dL) previously found to develop in apolipoprotein-E deficient mice by 10 weeks of age (27).

### Vasoreactivity Assays

Thoracic aortas were isolated from 7-mo (n=19/male; n=14/female) and 12-mo (n=15/male; n=18/female) C57BL/6J intact female (not cycled) and male mice, cut into 1 mm rings for isometric contractile force measurements. Thoracic aortas were placed in ice‐cold PSS bicarbonate buffer (PSS: 118.99mM NaCl, 4.69mM KCl, 1.17mM MgSO_4_*7H_2_O, 1.18mM KH_2_PO_4_, 1.60mM CaCl_2_-2H_2_O, 25.00mM NaHCO_3_, 0.03mM EDTA, 5.50mM glucose) bubbled continuously with 21% O_2_ and 5% CO_2_ and cleaned of fat and connective tissue and cut into 1mm long segments. Endothelium-intact or -denuded rings were mounted on tungsten wires and immersed in PSS at 37°C with constant gassing (21% O_2_ and 5% CO_2_) in a wire myography chamber (DMT 610) and incubated ex vivo with oxLDL (50*μ*g/dL; 2h; Kalen Biomedical, cat. no. 770202-7), followed by normalization and KCL (100mM) wakeup. Cumulative concentration-response curves to phenylephrine (PE) were generated to determine the contractile response. From these curves the EC50 was determined and used to contract rings to assess acetylcholine (ACh) dependent relaxation. BI-0115 (10*μ*M; selective LOX-1 inhibitor; Boehringer Ingelheim) was given 0.25h prior to oxLDL in some experiments to determine LOX-1 receptor dependence.

### Aortic Stiffness and Remodeling

Thoracic aortic stiffness was determined as previously described by assessing the slope of the stress-strain curve (28). In brief, diameter–tension relationships were determined following 2h incubation, but prior to KCL wakeup by a stepwise stretching of the tissue increasing its passive diameter by increasing the distance between the tungsten wires that were passed through the lumen of the aortic segment. Both the force and internal circumference of each aortic vessel segment was recorded and transformed into vessel diameter. The estimated diameter at 100mm Hg of pressure was determined utilizing the DMT normalization module (LabChart software, ADInstruments) which applies the obtained diameter-tension relationship and Laplace equation. Linear regression of each diameter-tension relationship was utilized to determine aortic stiffness and remodeling, corresponding to the steepness of the slope and diameter of vessel at zero pressure in relation to estimated diameter at 100mm Hg.

### Quantitative Real Time PCR

Quantitative real time PCR (qRT-PCR) was utilized to measure changes in mRNA levels of *LOX-1, CD36, ET-1, ET-1Rα, ET-1Rβ*, and *IL-6* in isolated 1 mo. and 16 mo. murine thoracic aortas as previously described (29). Primer sequences described in **Supplemental Table 1**.

### Reagents

All reagents were purchased from Sigma Aldrich (St. Louis, MO) unless otherwise noted.

### Data and Statistical Analysis

For each treatment group, an n≥3 was used to achieve an acceptable power for statistical analysis. Experiments were repeated for statistical analysis and data graphed using Prism 9.3.0 for Windows, GraphPad Software, USA, www.graphpad.com (GraphPad Software). Shapiro-Wilk and F-tests were performed to confirm if data sets achieved a normal distribution and equal variance among groups. Direct comparisons between two groups were made using an unpaired t test. Comparisons between three or more groups were made using a two-way ANOVA with Tukey’s multiple comparisons post hoc test. P < 0.05 was considered significant. Values are expressed as means ± SEM.

## RESULTS

### PE-induced contractile and ACh-induced relaxation responses are sex and age dependent in murine thoracic aortic ring segments

It has been previously shown that 26 mo. male C57BL/6 mice exhibited increased aortic contraction compared to younger 5 mo. mice in response to PE (30). Similarly, in this study thoracic aortic contraction was increased in both 12 mo. female and male mice relative to 7 mo. counterparts (**Figs. 1A-B**); however, this increase was most pronounced in the males (4.4x) as compared to the females (1.8x) (**Fig. 1B**). When comparing males to females there was a notable sex dependent increase in the efficacy of PE-induced contraction compared to males at 7 mo. that was not observed in 12 mo. mice (**Figs. 1C-D**). While there were notable effects in age and sex on PE-induced contraction efficacy we did not observe alterations to the potency of PE (**Fig. 1E**). From the PE cumulative concentration-response curves we derived the EC50 for each animal to next assess ACh-mediated relaxation. Counter to our observations of age-dependent differences in PE-induced aortic contraction, there were no differences in 7 and 12 mo. male and female ACh mediated relaxation (**Figs. 1F-G**). However, in alignment with previous observations made in 5 mo. male and female Sprague–Dawley rats (31) ACh-induced relaxation was greater in 7 mo. female mice than males (**Fig. 1H**). This sex difference in response to ACh was diminished in 12 mo. male and female mice (**Fig. 1I**). Similar to PE-induced contraction of the thoracic aortas, there was no change in the potency of ACh-mediated relaxation in the 7 mo. and 12 mo. males and female mice (**Fig. 1J**).

**Figure 1.**
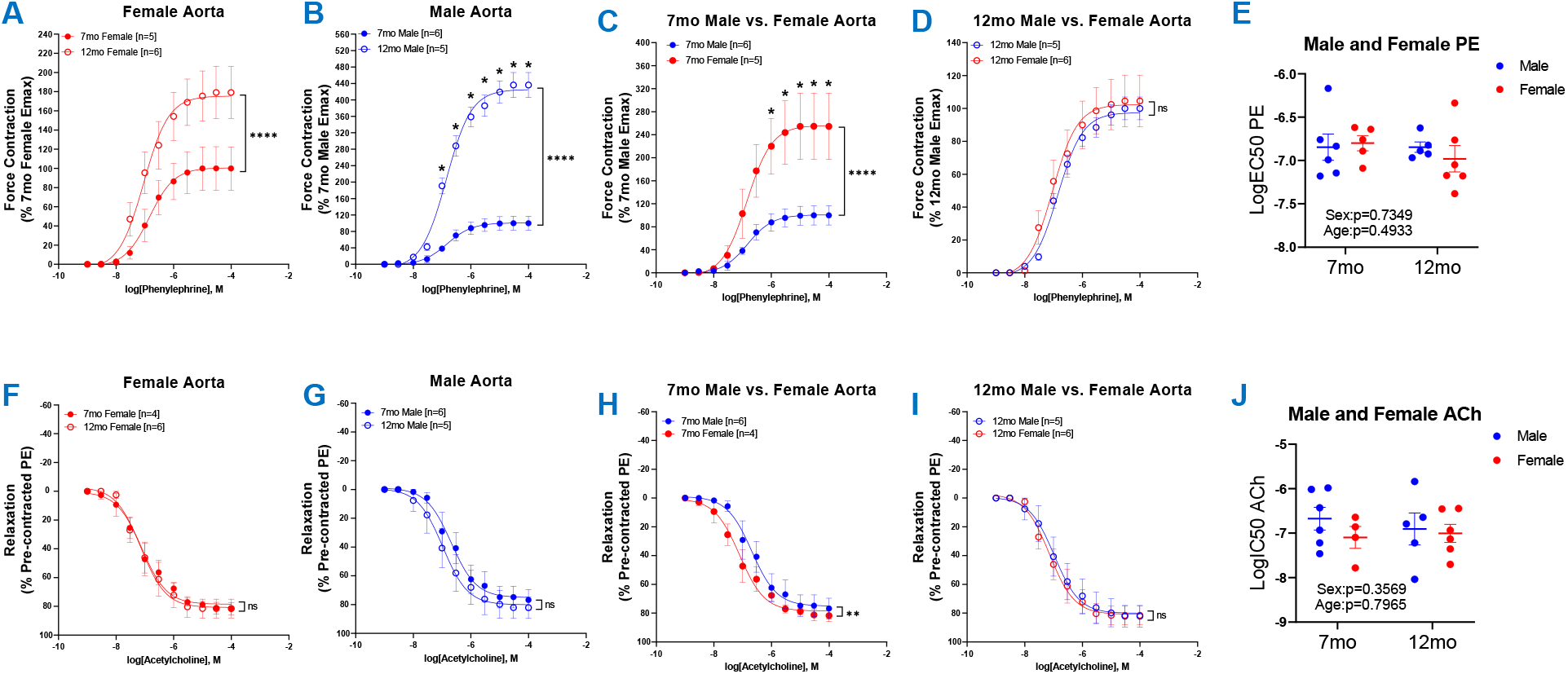
Vasoreactivity of 7 mo. and 12 mo. Male and Female Thoracic Aortic Rings. Concentration response curves to (**A-D**) PE and (**F-I**) ACh (PE-precontracted) in 1mm thoracic aortic rings from 7 mo. and 12 mo. male and female mice. (**E**) Graph representing LogEC50 of PE obtained from concentration response curves in male 7 mo. (n=6) and 12 mo. (n=5) and female 7 mo. (n=4-5) and 12 mo. (n=6) mice. (**J**) Graph represents LogIC50 of ACh obtained from concentration response curves in 7 mo. and 12 mo. male and female mice. Individual values are shown are transparent data points. Grouped data are represented as means ± SEM. Two-Way ANOVA with Tukey’s post-hoc test. *p<0.05, **p<0.01, ****p<0.0001.

### OxLDL increased PE-induced contraction and attenuated ACh-induced relaxation in male thoracic aortic rings in a LOX-1, age, and endothelial dependent manner

We next assessed the impact of acute oxLDL (50μ g/dL; 2h) exposure on 7 mo. and 12 mo. male thoracic aortic contraction and relaxation secondary to PE and ACh respectively. We first observed a prominent increase in the efficacy of PE-induced contraction in thoracic aortas incubated with oxLDL for 2h compared to vehicle (**Fig. 2A**). This oxLDL-mediated increase in PE contraction was then attenuated with selective LOX-1 inhibition (BI-0115; 10μ M), suggesting that LOX-1 mediates the observed oxLDL increase in PE-induced 7 mo. male thoracic aortic contraction (**Fig. 2A**). Additionally, when examining denuded 7 mo. male rings (**Supplemental Fig. 1A**), we observed that the oxLDL-mediated increase in PE efficacy was abrogated, and that LOX-1 inhibition alone or in the presence of oxLDL significantly attenuated contraction of the aortic rings (**Supplemental Fig. 1B**). Intriguingly, upon examination of the oxLDL/LOX-1 axis within 12 mo. male thoracic aortic rings, we observed a significant decrease in the efficacy of PE-mediated contraction in vessels incubated with oxLDL, LOX-1 inhibition, and the combination of oxLDL plus LOX-1 inhibition (**Fig. 2B**). However, we did not observe a change in the potency of PE-mediated contraction regardless of age or treatment (**Fig. 2C**). Following assessment of contractility, we next assessed ACh-mediated relaxation in the 7 mo. and 12 mo. pre-contracted thoracic aortic rings to further determine the effects of oxLDL/LOX-1 on the endothelium. In both the 7 mo. and 12 mo. male thoracic aortic rings, oxLDL significantly attenuated endothelial ACh-mediated relaxation (**Figs. 2D-E**). This response was LOX-1 dependent in the 7 mo. mice, but not in the 12 mo. male aortic rings. In addition, selective LOX-1 inhibition alone or in combination with oxLDL further enhanced ACh-mediated relaxation of the 7 mo. male aortic rings (**Fig. 2D**), but in the 12 mo. male aorta had a similar effect to oxLDL (**Fig. 2E**). Although there were changes in the efficacy of ACh following oxLDL, LOX-1 inhibition, and oxLDL plus LOX-1 inhibition there was no change in the potency of ACh-mediated relaxation (**Fig. 2F**). These data suggest that 1) acute oxLDL exposure has an oppositional effect on the contractility of 7 mo. and 12 mo. male thoracic aortas in response to PE which is endothelial LOX-1 dependent in the 7 mo. but not the 12 mo. males; 2) oxLDL induces endothelial dysfunction in both ages, but is more pronounced and LOX-1 dependent in the 7 mo. males compared to the 12 mo. males; 3) LOX-1 inhibition alone or in the presence of oxLDL further enhances endothelial function in the 7 mo. male thoracic aortas.

**Figure 2.**
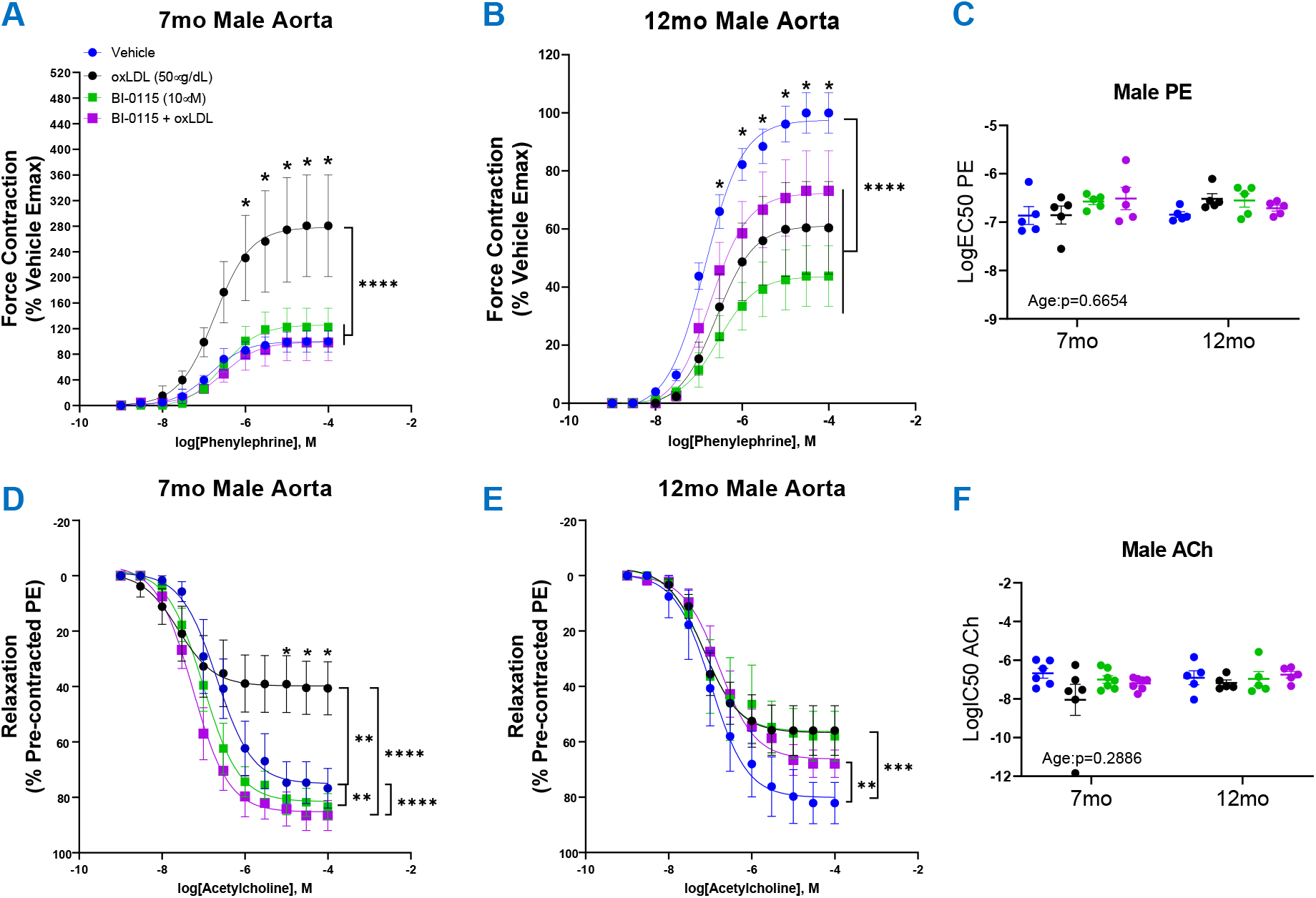
Vasoreactivity of 7 mo. and 12 mo. Male Thoracic Aortic Rings Following oxLDL Exposure, LOX-1 Inhibition, and oxLDL + LOX-1 Inhibition. Concentration response curves to (**A-B**) PE and (**D-E**) ACh (PE-precontracted) in 1mm thoracic aortic rings from 7 mo. and 12 mo. male mice following exposure to either vehicle (7 mo.:n=5-6; 12 mo.:n=5), oxLDL (50μ g/dL) (7 mo.:n=5-6; 12 mo.:n=5), BI-0115 (10μ M; selective LOX-1 inhibitor) (7 mo.:n=5-7; 12.mo.:n=5), or oxLDL (50μ g/dL) + BI-0115 (10μ M) (7 mo.:n=5-7; 12 mo.:n=5) for 2h. In aortas exposed to BI-0115, the selective inhibitor was administered 0.25h prior to oxLDL exposure. (**C**) Graph representing LogEC50 of PE obtained from concentration response curves in 7 mo. and 12 mo. male mice exposed as previously described. (**F**) Graph representing LogIC50 of ACh obtained from concentration response curves in 7 mo. and 12 mo. male mice exposed as previously described. Individual values are shown are transparent data points. Grouped data are represented as means ± SEM. Two-Way ANOVA with Tukey’s post-hoc test. *p<0.05, **p<0.01, ***p<0.001, ****p<0.0001.

### Female thoracic aortic ACh-induced relaxation was differentially altered a LOX-1, age, and endothelial dependent manner

It has been previously reported that LOX-1 expression was elevated in the aortas of C57BL/6J male mice compared to female mice (26) from which we hypothesized that oxLDL would differentially impact female aortic vasoreactivity compared to the males. We first assessed thoracic aortic contractility via PE in both 7 mo. and 12 mo. females and observed a decrease in contractility of both 7 mo. and 12 mo. females following exposure to either oxLDL, LOX-1 inhibition, or oxLDL plus LOX-1 inhibition (**Figs. 3A-B**). These results resemble our prior observation made in the 12 mo. male thoracic aortas; however, are in opposition to the contractility seen in the 7 mo. male aortas. Suggesting that the 12 mo. males may respond similarly to oxLDL and/or LOX-1 inhibition compared to females regardless of age. However, just as observed previously in the denuded male aortic rings, the effect of oxLDL was nullified in denuded female aortic rings but the lack of endothelium (**Supplemental Fig. 1C-D**) did not attenuate the effect of LOX-1 inhibition in the presence or absence of oxLDL (**Supplemental Fig. 1D**). When assessing the impact of oxLDL on endothelial dependent relaxation, we observed that oxLDL attenuated endothelial function in both 7 mo. and 12 mo. female aortic rings (**Figs. 3D-E**). These responses trended to be less severe in both the 7 mo. (51.5±9.1% relaxation) and 12 mo. (55.8±3.7% relaxation) female aortas than that observed in the 7 mo. male aortas (40.6±9.5% relaxation), but like the 12 mo. males (55.94±9.0% relaxation).

**Figure 3.**
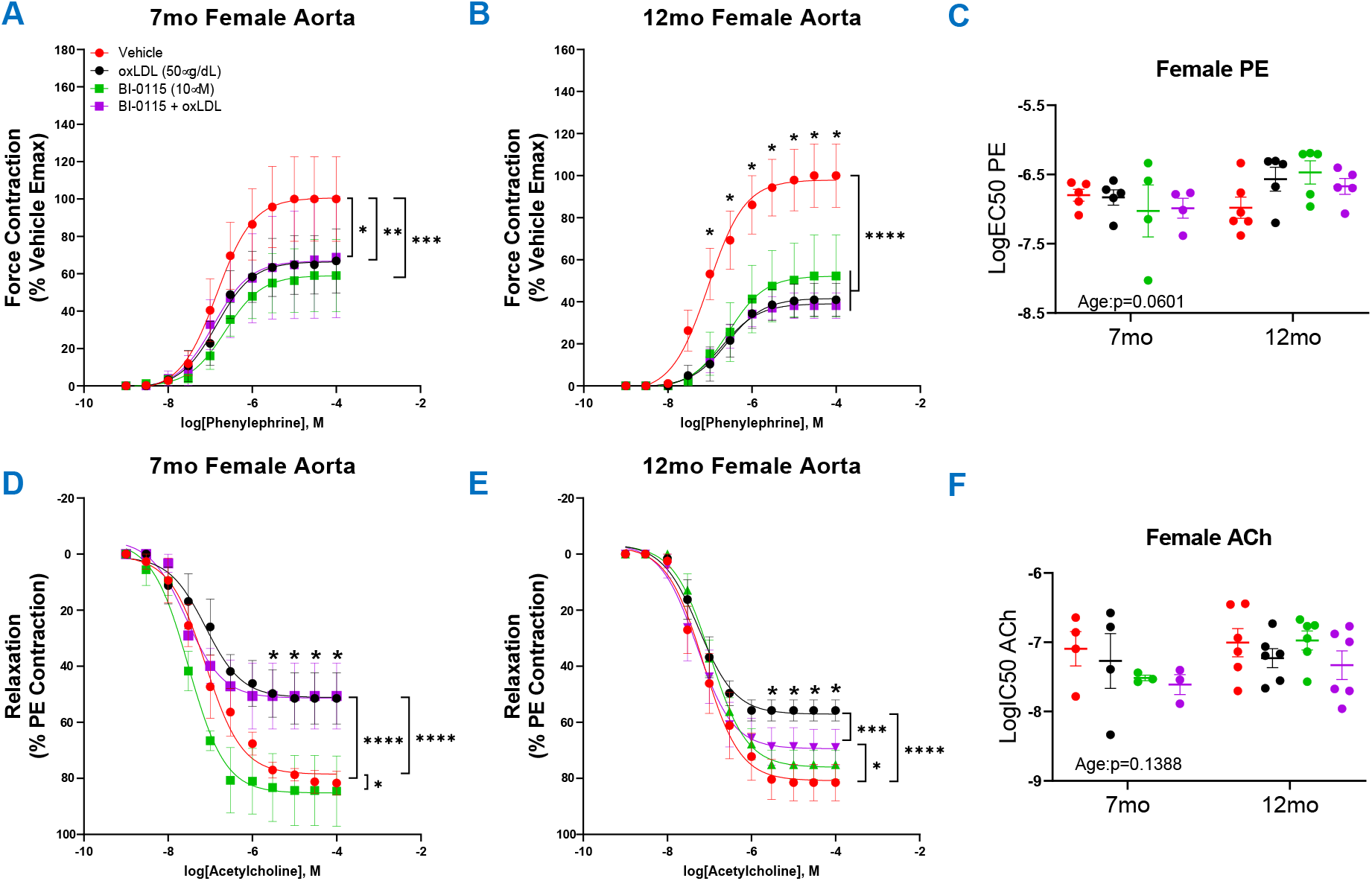
Vasoreactivity of 7 mo. and 12 mo. Female Thoracic Aortic Rings Following oxLDL Exposure, LOX-1 Inhibition, and oxLDL + LOX-1 Inhibition. Concentration response curves to (**A-B**) PE and (**D-E**) ACh (PE-precontracted) in 1mm thoracic aortic rings from 7 mo. and 12 mo. female mice following exposure to either vehicle (7 mo.:n=4-5; 12 mo.:n=5), oxLDL (50μ g/dL) (7 mo.:n=4-5; 12 mo.:n=6), BI-0115 (10μ M; selective LOX-1 inhibitor) (7 mo.:n=3-4; 12 mo.:n=5-6), or oxLDL (50μ g/dL) + BI-0115 (10μ M) (7 mo.:n=3-4; 12 mo.:n=5-6) for 2h. In aortas exposed to BI-0115, the selective inhibitor was administered 0.25h prior to oxLDL exposure. (**C**) Graph representing LogEC50 of PE obtained from concentration response curves in 7 mo. and 12 mo. male mice exposed as previously described. (**F**) Graph representing LogIC50 of ACh obtained from concentration response curves in 7 mo. and 12 mo. male mice exposed as previously described. Individual values are shown are transparent data points. Grouped data are represented as means ± SEM. Two-Way ANOVA with Tukey’s post-hoc test. *p<0.05, **p<0.01, ***p<0.001, ****p<0.0001.

Interestingly, just as observed in the 7 mo. male aortic rings, LOX-1 inhibition alone further enhanced endothelial function in the 7 mo. but not 12 mo. females (**Figs. 3D-E**). Moreover, oxLDL-mediated endothelial dysfunction in the 12 mo. females was attenuated by LOX-1 inhibition (**Fig. 3E**), but not in the 7 mo. females (**Fig. 3D**) which opposes the observations observed in the males (**Figs. 2D-E**). As observed previously, while oxLDL, LOX-1 inhibition, and oxLDL plus LOX-1 inhibition altered ACh efficacy, there was no change in potency of ACh (**Fig. 3F**). These data suggest that 1) acute oxLDL exposure decreases contractility of 7 mo. and 12 mo. female thoracic aortas in response to PE which is endothelial dependent and LOX-1 independent; 2) oxLDL induces endothelial dysfunction in both ages, and is partially LOX-1 dependent in the 12 mo. females but not the 7 mo. females; 3) LOX-1 inhibition alone further enhances endothelial function in the 7 mo. female thoracic aortas similar to that observed in the age matched males.

### OxLDL increases thoracic aortic stiffness and inward remodeling in an age, sex, endothelial, and LOX-1 dependent manner

We next examined how acute oxLDL exposure impacts the thoracic aorta functionally in terms of stiffness and remodeling as well as production of vasoactive and pro-inflammatory mediators. Utilizing the diameter-tension curves from both 7 mo. and 12 mo. male and female aortic rings we observed an overall effect of both sex and age on stiffness (**Fig. 4A**). Indicating that 7 mo. thoracic aortic rings were stiffer in comparison to 12 mo. mice and that male mice had increased stiffness compared to females. OxLDL as well as the selective LOX-1 inhibitor in the presence or absence of oxLDL increased stiffness in 7 mo. male aortas (**Fig. 4A**) in an endothelial dependent manner (**Supplemental Fig. 2A**). Whereas oxLDL did not influence the stiffness of 12 mo. males, or 7 mo. and 12 mo. females. Interestingly, the selective LOX-1 inhibitor alone or in the presence of oxLDL decreased 7 mo. female aortic stiffness (**Fig. 4A**) and was endothelial dependent (**Supplemental Fig. 2B**). While there was an overall effect of age on aortic stiffness, we did not observe an effect of oxLDL, LOX-1 inhibition, or oxLDL plus LOX-1 inhibition on 12 mo. male and female aortic stiffness. This contrasts our observations of remodeling as calculated by the vessel diameter at 100mm Hg of pressure previously shown to indicate inward remodeling (28). There was no overall effect of age on inward remodeling; however, we did observe a significant impact of sex suggesting that female thoracic aortas are smaller in diameter compared to males (**Fig. 4B**). Upon examination of the effects of oxLDL, dissimilar to our observations of altered stiffness we did not observe an effect of oxLDL on 7 mo. male and female remodeling (**Fig. 4B**). However, we did observe that in 7 mo. male denuded vessels, oxLDL, LOX-1 inhibition, and oxLDL plus LOX-1 inhibition increased inward remodeling (**Supplemental Fig. 2C**). Moreover, we observed that lack of endothelium resulted in inward remodeling of 7 mo. female aortic rings (**Supplemental Fig. 2D**). In contrast oxLDL induced a significant inward remodeling of intact 12 mo. male and female aortic rings that was LOX-1 dependent (**Fig. 4B**). From this we hypothesized that the 12 mo. mice may be potentially up taking oxLDL at a greater rate compared to the 7 mo. mice resulting in a smaller diameter at 100mmHg. In efforts to elucidate potential molecular mechanisms of oxLDL mediated altered vasoreactivity, stiffness, and remodeling we assessed levels of possible mediators within the thoracic aortas. We observed that oxLDL increased mRNA levels of LOX-1, endothelin-1, endothelin-1-receptors alpha and beta, and interleukin-6 (**Supplemental Fig. 3**). Intriguingly however, we did not observe an increase in CD36 expression following oxLDL exposure but did observe an increase with aging (**Supplemental Fig. 3**). Moreover, we also observed that levels of LOX-1, endothelin-1-receptor beta, and interleukin-6 were increased with age, but not endothelin-1 and endothelin-1-receptor alpha (**Supplemental Fig. 3**). Together these data suggest that increased age may potentiate a preferential increase in both LOX-1 and CD36-mediated uptake of oxLDL, resulting increased inward remodeling in 12 mo. mice.

**Figure 4.**
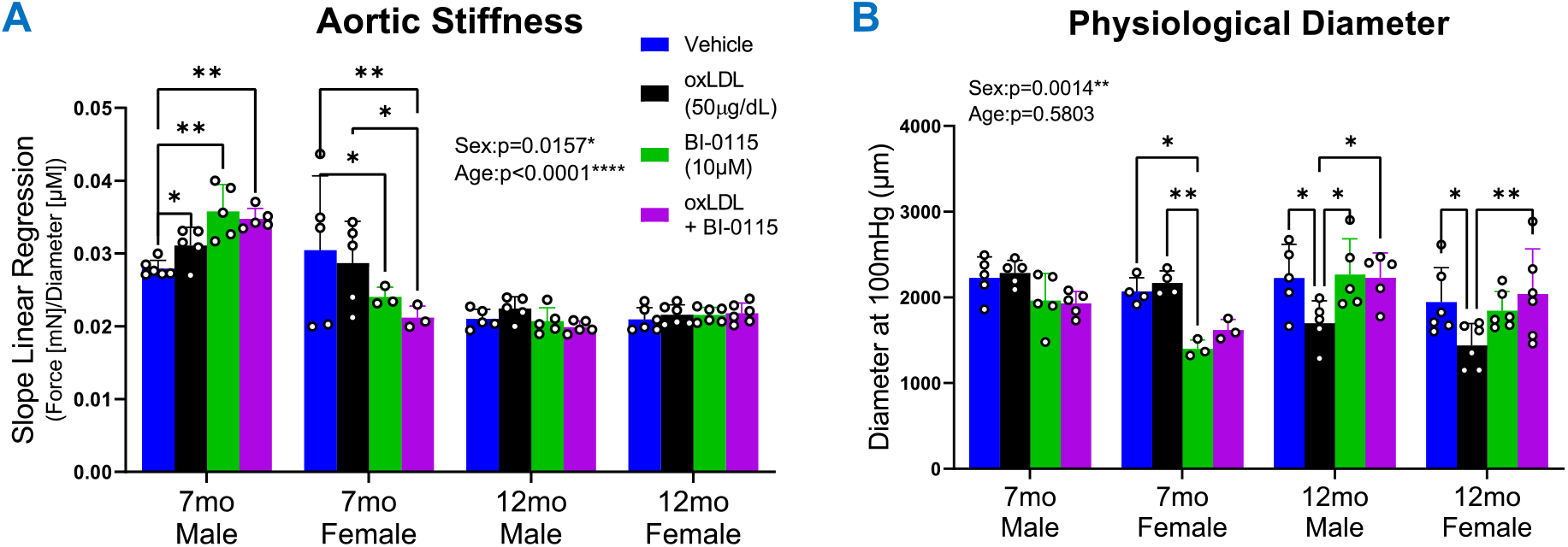
oxLDL Differentially Alters Thoracic Aortic Stiffness, and Remodeling. Graphs depicting wire myography mediated (**A**) linear regression analysis of diameter–tension relationships and (**B**) estimated diameter at 100mmHg from 1mm thoracic aortic vessels obtained from 7 mo. and 12 mo. male and female mice following exposure to either vehicle (Male:7 mo.:n=5; 12 mo.:n=5; Female: 7 mo.:n=4-5; 12 mo.:n=6), oxLDL (50μ g/dL) (Male:7 mo.:n=5; 12 mo.:n=5; Female: 7 mo.:n=4-5; 12 mo.:n=6), BI-0115 (10μ M; selective LOX-1 inhibitor) (Male:7 mo.:n=5; 12 mo.:n=5; Female: 7 mo.:n=3; 12 mo.:n=6), or oxLDL (50μ g/dL) + BI-0115 (10μ M) (Male:7 mo.:n=5; 12 mo.:n=5; Female: 7 mo.:n=3; 12 mo.:n=6) for 2h.

## DISCUSSION

In the present study we employed an ex vivo model to evaluate the impact of the oxLDL/LOX-1 axis on thoracic aortic vasoreactivity, stiffness, remodeling, and transcription in the context of sex and age to further elucidate the detrimental role oxLDL/LOX-1 plays in altering vascular function. Additionally, we evaluated the role of LOX-1 inhibition on the thoracic aortic vasculature and the role that the endothelium plays in regulating vascular function. To the best of our knowledge we have, for the first time, demonstrated the influence of sex, age, endothelium, and LOX-1 on acute ex vivo oxLDL exposure mediated altered aortic vasoreactivity, stiffness, and remodeling. In brief, oxLDL 1) increased contractility of 7 mo. male thoracic aortas in response to PE in an endothelial LOX-1 dependent manner, 2) decreased contractility in 12 mo. males as well as 7 mo. and 12 mo. females, 3) induced endothelial dysfunction in all animals regardless of age or sex, but was most pronounced and LOX-1 dependent in the 7 mo. males compared to 12 mo. males and 7 mo. and 12 mo. females, 4) increased aortic stiffness in 7 mo. males in a LOX-1 independent manner, and 5) increased inward remodeling of 12 mo. male and female thoracic aortas in a LOX-1 dependent manner.

In this study we observed that oxLDL exposure induced endothelial dysfunction resulting in endothelial dependent differentially altered vasoreactivity across age and sexes which appear to be in part LOX-1 receptor dependent. It has been previously found that oxLDL mediates an upregulation of endothelin-1, a potent vasoconstrictor, in endothelial cells (32), which corresponds to our findings at the mRNA and could in part play a role in our observations of increased vasocontractility in 7 mo. males. This is further perpetuated by findings demonstrating that blockade of ET-1R*α/β* improved endothelial-dependent vasodilation (33), which were observed to increase following oxLDL exposure in this study. While ET-1 is known to play a role in contractility, a separate study demonstrated that ET-1 is a potent vasoconstrictor in the abdominal aorta; however, in the thoracic aorta a 10nM dose of ET-1 only induced a 7.8% contraction compared to 60mM of K^+^ (34). Intriguingly, we observed a concomitant increase in basal expression of ET-1R*β* in 16 mo. male thoracic aortas but not alpha. Potentially suggesting an increase in ET-1R*β* mediated increase within the endothelium and subsequential increase in ET-1R*β*-mediated relaxation in the 12 mo. male mice resulting in attenuated contractility in response to PE as well as greater ACh-mediated relaxation, potentially due to previously demonstrated expression of ET-1R*β* within the endothelium (35-37) and an ET-1R*β*-mediated relaxation in rat thoracic aortic vessels (38). Expression of endothelin-1 and its receptors merits further investigation across the sexes as it has been previously observed that in human internal mammary arteries, exposure to 17β-estradiol resulted in downregulation of ET-1R*α/β* expression (39) suggesting that females may respond differentially along the endothelin-1/receptor axis compared to males and may in part delineate the decreased contractility observed in 7 mo. and 12 mo. female aortic rings exposed to oxLDL.

In addition to increased interest regarding vasoreactivity, there has been a burgeoning interest in elucidating the relationship between oxLDL and aortic stiffness as well as remodeling. Previous work such as the pivotal “Health ABC Study” (40), has contributed substantially to this line of interest. Wherein the authors explored the correlation between plasma oxLDL levels and arterial stiffness in older adults, revealing via pulse wave velocity measurements a significantly increased incidence of high arterial stiffness with increased oxLDL levels. Moreover, in a separate study, it was demonstrated that LOX-1 was associated with arterial stiffness in both middle-aged and elderly male and females (41). Intriguingly, in this study we observed a paradoxical decrease in aortic stiffness in the 12 mo. compared to 7 mo. mice. Counter to the previous report which demonstrated that male C57BL/6 mice thoracic aortas progressively increase in stiffness at 12 mo. compared to 4mo (30). We hypothesize that our observations could be due to a retention of elastic properties as we did not observe an age dependent change in luminal diameter as these have been described as the most consistent well-reported changes resulting in aortic stiffness (42). Further investigation into the underlying mechanisms of this observation is warranted but falls outside the scope of this study. While we observed no effect of oxLDL in 12 mo. male or female mice we did observe an increase in 7 mo. male thoracic aortic stiffness following oxLDL. Moreover, we observed a sex-dependent response of LOX-1 inhibition which resulted in increased 7 mo. male stiffness but decreased age-matched female aortic stiffness in the presence or absence of oxLDL in an endothelial dependent manner. Together suggesting that endothelial LOX-1 may play a differential role in the maintenance of 7 mo. male and female thoracic aortic stiffness but diminishes with age which could be a function of pathophysiologic aging-mediated upregulation of CD36 expression as observed in this study but warrants further elucidation. It has been reported that CD36 plays a critical role in oxLDL accumulation and internalization in macrophages (43, 44) from which we hypothesize that the upregulation of CD36 with age could also in part be responsible for the age dependent effects of oxLDL/LOX-1-mediated increases inward remodeling observed in the 12 mo. male and females, but not 7 mo. mice. Moreover, due to the previously noted LOX-1-mediated vascular remodeling (45), we theorize that CD36 is competing for oxLDL binding resulting in increased efficacy of LOX-1 inhibition to attenuate oxLDL/LOX-1-mediated inward remodeling in 12 mo. mice compared to 7 mo. mice. Together suggesting a differential role of LOX-1 in the progression of oxLDL-mediated thoracic aortic pathology in terms of stiffness and remodeling within males and female mice across age.

We acknowledge that the complex cascade of oxLDL/LOX-1 within the thoracic aorta cannot be exactly modeled in an ex vivo setting and that aortic rings do not exactly mimic the intact, pulsatile characteristics found in vivo. However, ex vitro studies do allow for the investigation of specific basic cellular and molecular mechanisms under conditions of oxLDL exposure which reflects what is observed during pathology such as hyper- and dyslipidemia.

### Perspectives and Significance

In conclusion, our ex vivo study has significantly contributed to the further elucidation of the intricate relationship between the oxLDL/LOX-1 axis and mouse thoracic aortic physiology, particularly within the contexts of sex and age. We have successfully demonstrated how acute ex vivo oxLDL exposure leads to distinct alterations in aortic vasoreactivity, stiffness, and remodeling, revealing differential responses across age and sexes. Importantly, our findings highlight the pivotal role of the endothelium and LOX-1 in coordinating these responses. Our results also further elucidate the interplay between oxLDL and endothelin-1, contextualizing their potential contributions to observed vasoreactivity changes. Furthermore, our paradoxical observation of decreased aortic stiffness in older mice warrants deeper exploration to uncover the underlying mechanisms. While we observed varying effects of LOX-1 inhibition on stiffness between age-matched male and female mice, its potential relationship with CD36-mediated processes adds yet another layer of complexity to the complex nature of oxLDL-mediated effects. Future studies will be aimed at further elucidation into the mechanisms linking oxLDL, LOX-1, endothelin-1, and CD36, while also considering the broader implications for age and sex differences in vascular health. This study lays a strong foundation for unraveling the elaborate molecular pathways that contribute to oxLDL/LOX-1-mediated alterations in aortic physiology, with potential implications for therapeutic interventions in cardiovascular health.

## DATA AVAILABILITY

All materials and data will be available from the corresponding author following University of Arizona’s policy of sharing research materials and data.

## ACKNOWLEDGMENTS

We would like to thank Dr. Taben Hale for providing us access to and utilization of the DMT 610 myography rig.

## DISCLOSURES & DISCLAIMERS

The authors declare no conflict of interest.

## AUTHOR CONTRIBUTIONS

Conceptualization, T.S.W and R.J.G.; methodology, T.S.W and R.J.G.; software, T.S.W and R.J.G.; validation, T.S.W and R.J.G.; formal analysis, T.S.W and R.J.G.; investigation, T.S.W and R.J.G.; resources, T.S.W and R.J.G.; data curation, T.S.W and R.J.G.; writing-original draft preparation, T.S.W.; writing-review and editing, T.S.W and R.J.G.; visualization, T.S.W and R.J.G.; supervision, R.J.G.; project administration, R.J.G.; funding acquisition, T.S.W and R.J.G. All authors have read and agreed to the published version of the manuscript.

## SUPPLEMENTAL FIGURE LEGENDS

**Supplemental Figure 1. Vasoreactivity of 7 mo. Male and Female Endothelial Denuded Thoracic Aortic Rings**. Concentration response curves to (**B&D**) PE and (**A&C**) ACh (PE-precontracted) in 1mm denuded thoracic aortic rings from 7 mo. male and female mice following exposure to either vehicle (Male: n=3; Female: n=4), oxLDL (50*μ*g/dL) (Male: n=3; Female: n=4), BI-0115 (10*μ*M; selective LOX-1 inhibitor) (Male: n=5; Female: n=3), or oxLDL (50*μ*g/dL) + BI-0115 (10*μ*M) (Male: n=5; Female: n=3) for 2h. Individual values are shown are transparent data points. Grouped data are represented as means ± SEM. Two-Way ANOVA with Tukey’s post-hoc test. *p<0.05, ****p<0.0001.

**Supplemental Figure 2. Aortic Stiffness and Remodeling of 7 mo. Male and Female Endothelial Intact and Denuded Thoracic Aortic Rings**. Graphs depicting wire myography mediated (**A-B**) linear regression analysis of diameter–tension relationships and (**C-D**) estimated diameter at 100mmHg from 1mm thoracic aortic vessels obtained from 7 mo. and 12 mo. endothelial intact and denuded male and female mice following exposure to either vehicle either vehicle (Male: n=4-5; Female: n=4-5), oxLDL (50μ g/dL) (Male: n=4-5; Female: n=4-5), BI-0115 (10μ M; selective LOX-1 inhibitor) (Male: n=5; Female: n=3-4), or oxLDL (50μ g/dL) + BI-0115 (10μ M) (Male: n=5; Female: n=3-4) for 2h. Data are represented as means ± SEM. Two-Way ANOVA with Tukey’s post-hoc test. *p<0.05, **p<0.01, ***p<0.001, ****p<0.0001.

**Supplemental Figure 3. oxLDL Differentially Alters Thoracic Aortic Stiffness, Remodeling, and Vasoreactivity and Pro-Inflammatory Mediators**. qRT-PCR graph of thoracic aortas from 1 mo. male exposed to either vehicle (n=4) or oxLDL (50μg/dL; n=5) and 16 mo. male mice exposed to vehicle (n=3-4) for 2h. Data are represented as means ± SEM. Two-Way ANOVA with Tukey’s post-hoc test. *p<0.05, **p<0.01, ***p<0.001, ****p<0.0001.

**Supplemental Table 1. qRT-PCR Primers for LOX-1, CD36, ET-1, ET-1Ra, ET-1Rb, and IL-6**.

