## Supplementary figures and images for "OxLDL/LOX-1 Mediated Sex, Age, and Cell Dependent Alterations in Mouse Thoracic Aortic Vascular Reactivity"

### Supplemental Figure 1

#### Supplemental Figure 1

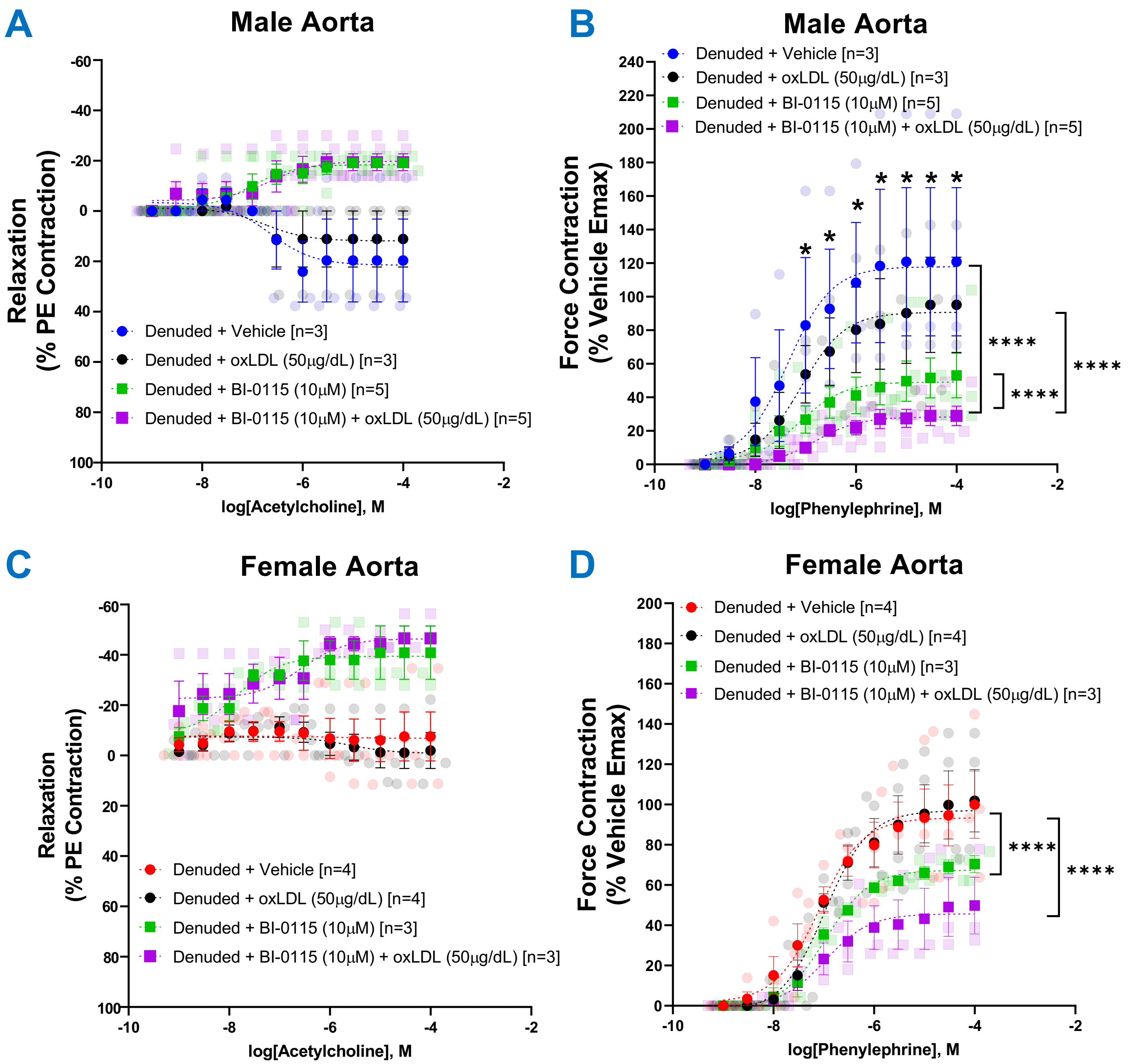

### Supplemental Figure 2

#### Supplemental Figure 2

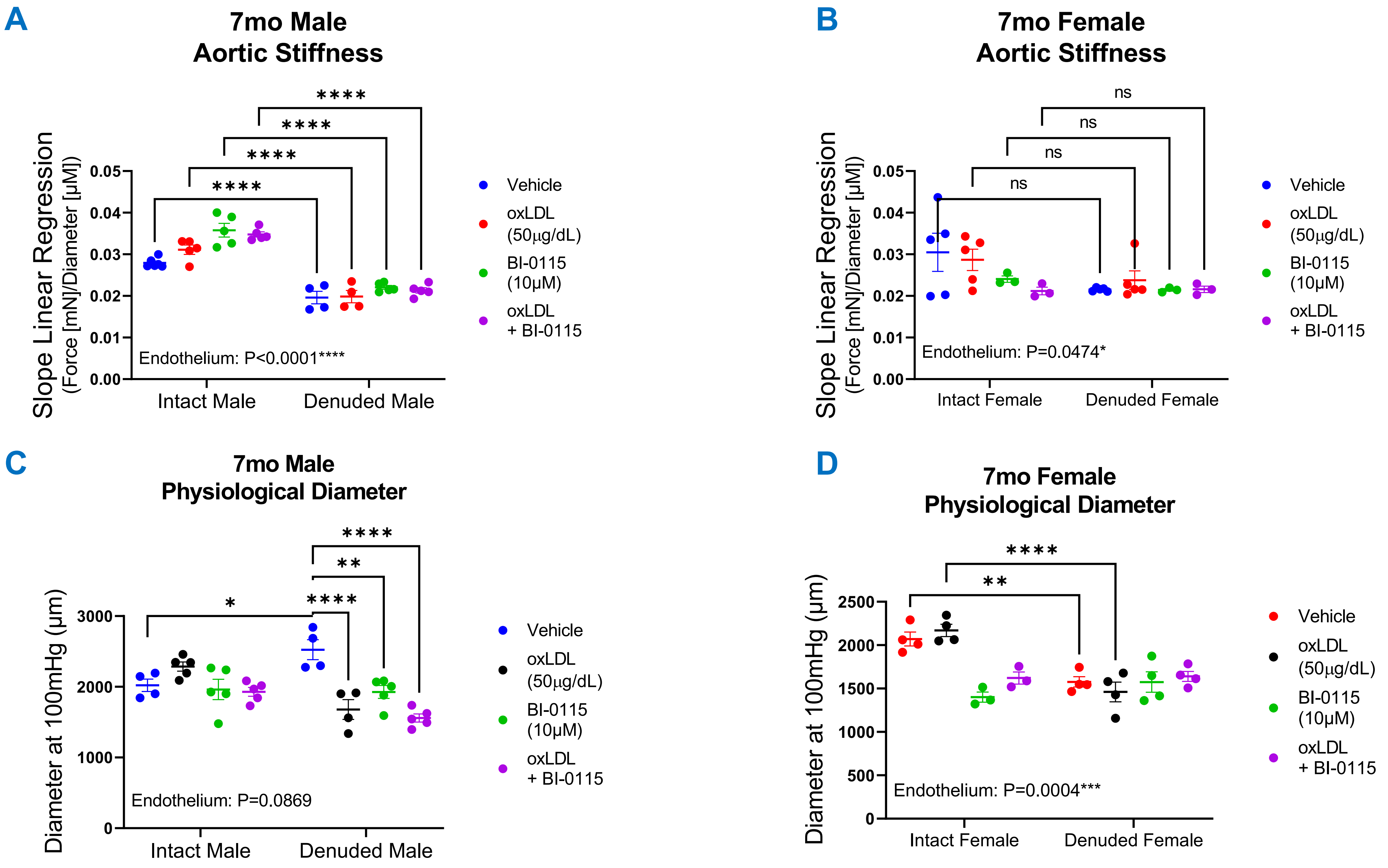

### Supplemental Figure 3

**Supplemental Figure 3**

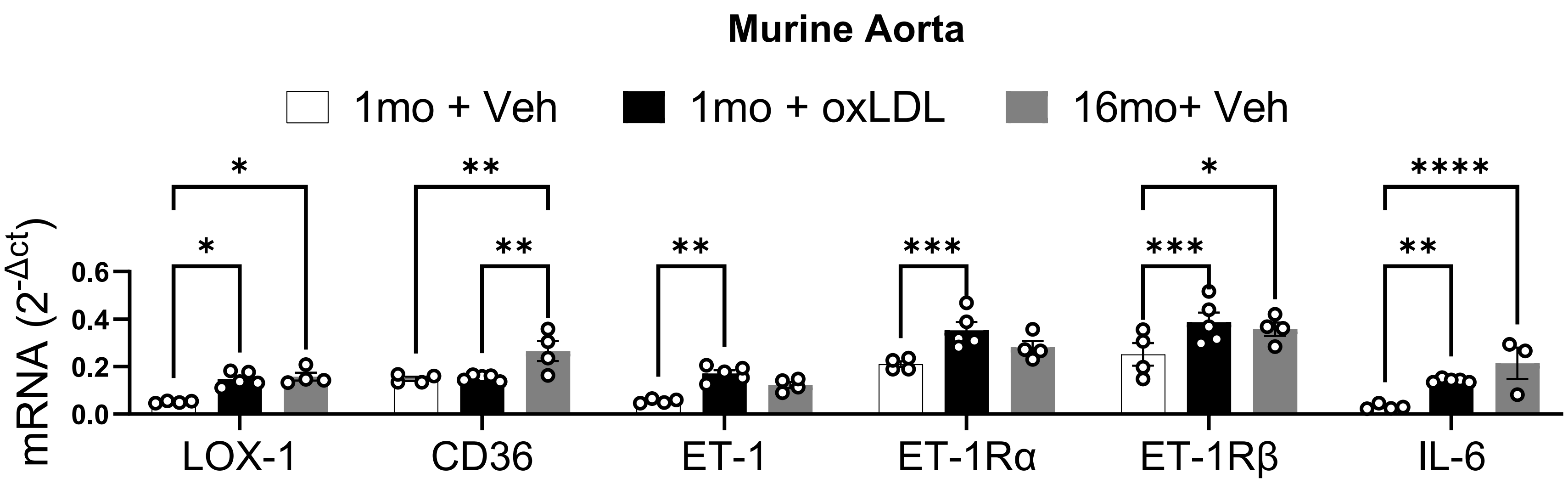
