## Supplemental Table 1 for "OxLDL/LOX-1 Mediated Sex, Age, and Cell Dependent Alterations in Mouse Thoracic Aortic Vascular Reactivity"

| Primer | Sense (5'-3') | Anti-Sense (5'-3') |
|---|---|---|
| **LOX-1** | GCTCTGCTTCTCGTGGGCAT | CGAAGGCCCCAAGGAAAGGG |
| **CD36** | TGTGGCAAACAGGGCTGGAG | GCAAGCACAAGTCTGGATCACC |
| **ET-1** | ACGCCAGTGCTAATGGCTCC | AGGTGTCTGCACTCAAGGCG |
| **ET-1Ra** | TGCTTTGATCAGGCACCCTCC | CCCAGAGCTGACTTCTGCCG |
| **ET-1Rb** | ATGCCCTGATGCCTTAGCCAC | ACCCACCTGCAGAGCAAGAAC |
| **IL-6** | CCAAGAGGTGAGTGCTTCCCC | ACTCTCTCCCTTCTGAGCAGC |
